# Repopulation of a 3D-simulated periapical lesion cavity with triggered osteoblastic-differentiated dental pulp stem cell spheroids

**DOI:** 10.1101/2021.07.02.450930

**Authors:** Vítor Luís Ribeiro, Janaína A. Dernowsek, Roger R. Fernandes, Dimitrius L. Pitol, João Paulo Mardegan Issa, Jardel F. Mazzi-Chaves, Karina Fittipaldi Bombonato-Prado, Manoel Damião Sousa-Neto, Geraldo Aleixo Passos

## Abstract

We established a proof of concept model system for the biological repair of periapical lesions using stem cell spheroids. A mesenchymal stem cell line isolated from the dental pulp of deciduous teeth (shed cells) was cultured in a 2D monolayer and then in 3D multicellular spheroids. An image of a periapical lesion of an upper lateral incisor tooth was obtained by computed micro tomography, which was used as a model for photopolymer resin 3D printing to generate a negative frame of the lesion. The negative model served to prepare a positive model of the periapical lesion cavity in an agarose gel. Shed cells cultured in monolayers or as spheroids were seeded in the positive lesion mold before or after osteoblastic differentiation. The results showed that compared to cells cultured in monolayers, the spheroids featured uniform cellularity and had a greater viability within the lesion cavity, accompanied by a temporal reduction in the expression of mRNAs typically expressed by stem cells (Cd13, Cd29, Cd44, Cd73, and Cd90). Concomitantly, there was an increase in the expression of protein markers that characterize osteoblastic differentiation (RUNX2, ALP, and BGLAP). These results provide a new perspective for regenerative endodontics with the use of spheroids prepared with shed cells to repair periapical lesions.

## Introduction

Anaerobic polybacterial infection of the dental pulp and root canals causes inflammation and periapical injury. Inflammation may harm and damage tissues, resulting in reabsorption of the bone around the root apex (1). To repair the periapical lesion, intervention is necessary to eliminate or reduce the bacterial contingency present in the root canals. These goals are achieved through endodontic treatment, which consists initially of the biomechanical preparation of the root canal system, irrigating solutions to dissolve necrotic organic tissues, intracanal medications, and canal filling, followed by restorative processes. The treatment allows the restoration of the function of the dental element (2–4) and subsequent sealing so that microleakage or new bacterial contamination of the root canal system does not occur.

As periapical injury is a multifactorial disease, factors other than endodontic and restorative treatments are involved. These factors, such as the bacterial microbiology related to the intrinsic characteristics of the host, either by its ability to defend itself in response to the contaminating agent or its ability to repair after removal, need to be taken into account. Thus, the relationship between host resistance and the virulence of the microorganisms directly influences disease development and repair. In other words, not all individuals have the same phenotypic response and gene expression related to a given treatment (5–7).

Patients who do not have self-regenerating ability (8,9) could benefit from therapies based on stem cells, scaffolds, and growth and differentiation factors involved in dental pulp renewal. This strategy is promising, which might result in healing the disease, the longevity of dental elements and the regeneration of persistent or critical size periapical lesions (9–11).

The stem cells of human exfoliated primary teeth (shed cells) are mesenchymal stem cells obtained from dental pulp explants (12,13). Shed cells are being widely used in studies because they feature a high proliferation rate and express markers such as Oct4, CD13, CD29, CD44, CD73, CD90, CD105, CD146, and CD166 that are used for their characterization (14–16). These cells have been shown to be important in the regeneration and repair of craniofacial injuries, loss of teeth, and bones due to their high proliferation and differentiation capacity. They might be suitable for regenerative endodontic therapy for treatments involving the regeneration of critical-sized bone lesions and persistent periapical lesions (8,17,18).

Monolayer (2D) mammalian adherent cell cultures have been the gold-standard method for cell biology studies for many years. However, changes in the cell’s properties may occur, leading to differences with the organism’s *in vivo* microenvironment (19–21). Mammalian cells, in general, are very responsive. Depending on the physical or chemical stimulus, they can show adaptive changes in their biochemical or physiological mechanisms. In an attempt to reduce the effects of monolayer (2D) culture and get closer to the *in vivo* condition, three-dimensional (3D) cell cultures have been developed (22,-24). A 3D structure of cells can be achieved by forming nonadherent multicellular spheroids, which show a more remarkable similarity of biochemical and physiological responses to *in vivo* tissues than adherent cells cultured in 2D (25,26).

In this work, we established a model system using shed cells to form spheroids *in vitro* and compared them with shed cells cultured in 2D in regards to their osteoblastic differentiation capacity. In addition, we compared spheroids versus cells cultured in 2D during their osteoblastic differentiation within an actual 3D simulated periapical lesion model.

## Material and Methods

### Mesenchymal stem cells from human exfoliated deciduous teeth (shed cells)

Shed cells used in this study were previously isolated (27,28). The shed cells were separated from a normal male child and cultured in DMEM with HAM F12 supplemented with 15% fetal bovine serum (FBS) plus 100 U/ml penicillin and streptomycin (here referred to as control medium or CMD medium) at 37°C in an incubator with a 5% CO_2_ atmosphere. These cells were previously assayed by flow cytometry using a BD FACSCalibur flow cytometer (Becton Dickinson and Company, Franklin Lakes, NJ) to confirm the expression of the surface markers CD29, CD44, CD13, CD90, and CD73 (17). This cell line showed no contamination with hematopoietic stem cells, as determined by the assessment of the following surface markers: CD34 and CD45/CD14 (17).

In this study, the cells were cultured as both conventional adherent 2D monolayers and nonadherent 3D spheroids in Dulbecco modified Eagle Medium (DMEM) supplemented with nutrient mixture F-12 (DMEM/F12, Gibco^®^) with 15% inactivated fetal bovine serum in an incubator at 37°C with a 5% CO_2_ atmosphere.

All of the procedures involving the sampling of human cells were approved by the Committee for Ethics in Research of the Biosciences Institute, USP, São Paulo, Brazil (approval # 435054) for Dr. Maria Rita S. Passos-Bueno, who isolated the cells and established the cell line. She kindly supplied the shed cells from the original stock for this study.

### Formation of spheroids from the shed cells

In this study, microwells in a 2% agarose gel prepared with sterile PBS were used and molded on a set of 600 μm photopolymer resin pins.

The agarose microwells were filled with a suspension of trypsinized shed cells (1.25 × 10^5^ cells/ml DMEM/F12 medium) from conventional monolayer 2D cultures. The agarose microwells with the shed cells were incubated inside 12-well multiwell plates at 37°C and 5% CO_2_ for 24 to 96 hours to allow the formation of multicellular spheroids, whose formation was observed through a Cytosmart™ Live Cell Imaging System (Lonza).

### Simulation of a periapical lesion cavity by 3D printing

An *in vitro* model system to mimic a periapical lesion was used in this study. It was made by scanning the area of a periapical lesion of a tooth that needed endodontic treatment. This area was obtained using cone beam computed tomography of an upper lateral incisor tooth of a patient from the collection of tomographic examinations at the Endodontics Clinic at the School of Dentistry of Ribeirão Preto, University of São Paulo, Ribeirão Preto, SP, Brazil. The file was segmented using the CTAn program v.1.18.4.0+ (SkyScan, Kontich, Belgium) and using a histogram of contrasts. They were obtained through the differences in gray density, the segment, and the separation of the structure that makes up the periapical lesion of adjacent structures (tooth and alveolar bone).

The 3D periapical lesion molds were designed by reverse engineering using Rhinoceros^®^ 5.0 software (McNeel North America, Seattle, WA, USA), which generated a stereolithographic file (STL) of the scanned periapical lesion. The procedure was useful for the determination of a periapical lesion model with actual dimensions.

Subsequently, the molds were made using 3D printing. The STL file was executed using digital light processing (DLP) technology on a SparkMaker^®^ 3D printer (LCD-E, SparkMaker Corporation, Shenzhen, China) so that the negative molds were reproduced with the exact shapes and measurements of the periapical injury scanned. The 3D molds were designed to be used in 12-well multiwell culture plates, containing three periapical lesions of identical shapes and measurements per mold. The positive molds on agarose were carried out using the agarose gel preparation protocol as described above.

The positive molds simulated the actual periapical lesion, which was used for later seeding of shed cells under four different conditions: 1) shed cells from 2D cultures, multicellular 3D shed cell spheroids (from 600 μm pin micromolds), 3) shed cells from 2D cultures after osteoblastic differentiation, and 4) multicellular 3D shed cell spheroids after osteoblastic differentiation (from 600 μm pin micromolds).

### Construction of growth curves of spheroids

Two spheroid growth curves were built, one for each type of agarose micromold: a spheroid curve for those growing in 600 μm micromolds, and another for spheroids growing in micromolds mimicking the periapical lesion. For the growth curve in 600 μm microwells, 1.25 × 10^5^ cells were seeded, and the spheroids were dispersed by conventional trypsin treatment 24, 48, 72, and 96 hours after seeding the shed cells. For the growth curve of spheroids growing in the micromolds of the periapical lesion, 3.0 × 10^5^ cells were seeded, and the spheroids were dispersed with trypsin 8, 16, 24, 32, and 40 hours after seeding. The cells were stained with trypan blue and counted in an automatic cell counter (Nexcelom Auto T4 Cellometer). For each of the curves, the arithmetic mean of three determinations for each time point and the mean’s standard deviation were calculated.

### Spheroid histology

The spheroids formed in the 600 μm micromolds or the periapical lesion micromolds were harvested during the exponential growth phase. The spheroids were fixed in 10% pH 7.0 buffered formaldehyde and dehydrated in ethanol using a conventional hematoxylin-eosin (H&E) histology protocol. The dehydrated material was mounted on a microscopy slide, stained with H&E, and covered with a coverslip for later examination under an optical microscope (Leica, Germany).

### Viability assays

### Monolayer cell culture viability

Samples of shed cells cultured with DMEM/F12 medium plus chemical osteoblastic inducers were collected from three time points of differentiation, i.e., after 3, 7, and 10 days, for cell viability assessment using the CyQuant™ MTT Cell Viability Assay (Thermo Fisher), which quantitatively measures lactate dehydrogenase (LDH) released into the culture medium as a result of cell membrane damage and cell death, according to the manufacturer’s instructions.

### Spheroid viability

The qualitative viability of spheroids formed in 600 μm micromolds or in micromolds of the periapical lesion and at the exponential growth phase was evaluated using the Live/Dead™ Viability/Cytotoxicity Kit (Invitrogen-Thermo Fisher Scientific), according to the manufacturer’s instructions. The procedure discriminates live from dead cells based on staining with green-fluorescent calcein-AM as an indicator of intracellular esterase activity of live cells and red-fluorescent ethidium homodimer-1 as an indicator of loss of plasma membrane integrity of dead cells.

The images were recorded using a fluorescence microscope (Nikon model Ti 5). The regions of spheroids stained green represent living cells, and those stained red represent dead cells.

### In vitro osteoblastic differentiation of shed cells

The chemical induction of differentiation of shed cells, cultured in 2D monolayers, into osteoblasts was triggered by the addition of 10^−7^ M dexamethasone (Sigma), 5 μg/mL ascorbic acid (Gibco), and 2.16 g/mL β-glycerolphosphate (Sigma) to the DMEM/F12 culture medium supplemented with 15% FBS (17,30). During the entire culture time, the cells were kept at 37°C in an atmosphere with 5% CO_2_. The culture medium containing the chemical inducers was changed every three days.

Samples were collected from three time points of differentiation, i.e., after 7, 14, and 21 days, for cell viability and osteoblastic differentiation assays, such as MTT, Fast Red, and mineralization assays, immunofluorescent staining with antibodies recognizing osteoblastic cell markers and RT-qPCR to quantify the expression of mRNAs related to osteoblasts.

Additionally, at the 7- and 14-day time points at which the cells were in contact with the inducing agents, the cells were removed from the culture flasks by trypsinization and seeded in the 600 μm or in the periapical lesion agarose micromolds to obtain spheroids during the osteoblastic differentiation process.

The monitoring of osteoblastic differentiation was performed using the methods described below.

### Detection of alkaline phosphatase (ALP)

Briefly, shed cells were cultured in the presence or absence of inducing agents in 24-well plates and evaluated at time points of 7, 10, 14, and 21 days. ALP activity was detected by reaction with Fast Red (Sigma Aldrich) (31). After staining, the liquid medium was removed, and the plates were allowed to dry for 48 hours at room temperature and photographed. The images were analyzed with the ImageJ program available at (https://imagej.nih.gov/ij/download.html).

### Mineralization

Mineralization (calcium deposition areas) was evaluated by staining with Alizarin Red (32). Shed cells were grown in the presence or absence of inducing agents in 24-well plates and assessed at 7, 14, 17, and 21 days. The liquid medium was removed from the cultures, and the wells were washed three times with PBS at 37°C, dehydrated with 70% ETOH for one hour at 4°C, washed with PBS and stained with Alizarin Red (Sigma Aldrich) for 45 minutes. After this period, the wells were washed with PBS and deionized water and allowed to dry for 48 hours at room temperature. The wells were photographed, and the images were analyzed using the ImageJ program.

### Immunofluorescence staining

For immunofluorescence staining, we used a protocol previously described (33). Briefly, shed cells grown in a monolayer on glass coverslips in control medium or medium plus chemical inducers for 7, 14, and 21 days were fixed in 4% paraformaldehyde in PBS pH 7.2 for 10 minutes at room temperature.

Permeabilization was performed with 0.5% Triton X-100 solution in PBS for 10 minutes, followed by blocking with a 5% solution of skimmed milk in PBS for 30 minutes. Primary monoclonal antibodies against RUNX2 (IgG mouse anti-human monoclonal antibody from Abcam, UK, 1:100 dilution), ALP (IgG1 mouse anti-human B4-78 monoclonal antibody from Developmental Studies Hybridoma Bank, USA) (1:100 dilution), or BSP (IgG2a mouse anti-rat WV1D1 (9c5), a monoclonal antibody (Developmental Studies Hybridoma Bank, USA) (1:200 dilution) were incubated with the cells for 1 hour.

After incubation with primary antibodies, cells were stained with goat anti-mouse secondary antibody conjugated with Alexa Fluor 594 (Molecular Probes, Thermo Fisher Scientific, USA), 1:200 dilution, for 50 minutes. For the observation of cell limits and cell nuclei, phalloidin conjugated with Alexa Fluor 488 (Molecular Probes, USA) at a 1:200 dilution was used, and the cells were incubated for 50 minutes and DAPI (4′,6′-diamino-2-phenylindole) (Molecular Probes, Thermo Fisher Scientific, USA) for 5 minutes. Antifade Prolong mounting medium (Molecular Probes, USA) was used, and labeling was analyzed using a Leica fluorescence microscope model DMLB (Leica Microsystems, Wetzlar, Germany).

### Reverse transcription quantitative real-time PCR (RT-qPCR)

RT-qPCR was also used to evaluate the transcriptional expression levels of mRNAs involved in osteoblastic differentiation or those related to the stemness of mesenchymal cells. The mRNAs (cDNAs) are indicated as follows, with the respective GenBank accession numbers and sequences of their respective 5′ to 3′ forward and reverse oligonucleotide primers in parentheses: Gapdh acc, (ACGACCAAATCCGTTGACTC and CTCTGCTCCTCCTGTTCGAC); Runx2 acc, (AGTAAGAAGAGCCAGGCAGG and GCTGGATAGTGCATTCGTGG); Alpl acc, (CCACGTCTTCACATTTGGTG and AGACTGCGCCTGGTAGTTGT); Bglap acc, (GGCAGCGAGGTAGTGAAGAG and CTGGAGAGGAGCAGAACTGG); Sp7 acc, (TGCTTGAGGAGGAAGTTCAC and AGGTCACTGCCCACAGAGTA); Bsp acc, (ACAACACTGGGCTATGGAGA and CCTTGTTCGTTTTCATCCAC); Cd13 acc, (GCCGTGTGCACAATCATCGC and CACCAGGGAGCCCTTGAGGT); Cd29 acc, (ACCAAGGTAGAAAGTCGGGA and TGACCACAGTTGTTACGGCA); Cd44 acc, (ACTGCAATGCAAACTGCAAG and AAGGTGGAGCAAACACAACC); Cd73 acc, (GTTCTCCCAGGTAATTGTGC and ACCTGAGACACACGGATGAA); Cd90 acc, (GCCCTCACACTTGACCAGTT and GCCTTCACTAGCAAGGACGA).

Thermal cycling was completed using a StepOne Real-Time PCR System (Applied Biosystems) as follows: 50°C for 2 min, 95°C for 15 min, and 60°C for 1 min (40 cycles). The 2^−ΔΔCT^ method was used as a relative normalization method. The Gapdh mRNA expression level was used as an internal housekeeping normalizer. We used the GraphPad Prism 5.00 tool (http://www.graphpad.com/prism/Prism.html) to run one-way ANOVA with Bonferroni’s correction to compare the expression levels of stemness mesenchymal mRNAs (Cd13, Cd29, Cd44, Cd73 and Cd90). To compare the expression levels of osteoblastic differentiation mRNAs (Runx2, Alpl, Bglap, Sp7 and Bsp), unpaired t-tests were used. For both types of statistical tests, p < 0.05 was considered significant.

## Results

### Spheroid morphology and growth curves

Spheroid morphology was determined by optical microscopy of histological sections stained with H&E. Fig. 1 shows histological sections of the three types of spheroids formed according to the inoculum of cells and agarose molds.

**Figure 1.**
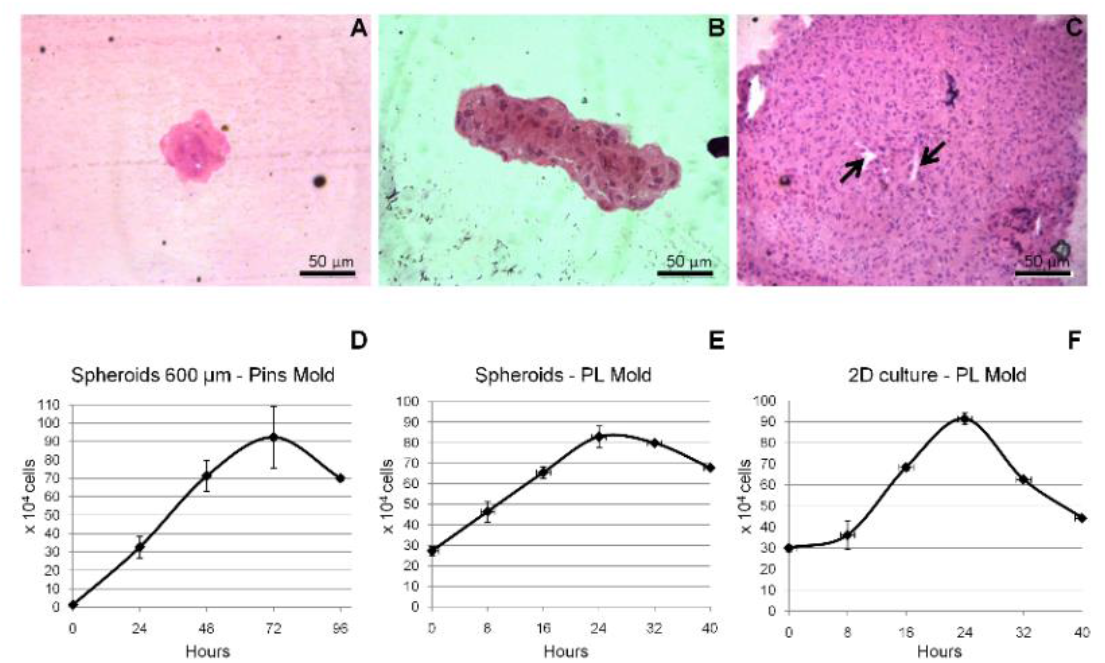
Spheroid morphology and growth. Spheroids were grown in three different microenvironments and then processed for histological examination. In Fig. 1A, it is possible to observe a typical type “A” spheroid grown for 24 hours in a 600 μm agarose mold. This type of spheroid assumed spherical morphology with well-grouped cells. In Fig. 1B, it is possible to observe a typical type “B” spheroid grown in a 600 μm agarose mold for 24 hours and then transferred to a periapical lesion mold. This type of spheroid presented a compact and spindle-shaped morphology. In Fig. 1C, it is possible to observe a typical type “C” spheroid, which was grown from cells from 2D monolayer culture transferred to a periapical lesion mold. The type “C” spheroid thus grown showed a spherical morphology comparable to the type “A” spheroids but larger and with a hollow nucleus without cells (arrows). The three types of spheroid’s growth curves presented the growth, stationary, and decline phases (Figs. 1D-F). Time-point values are presented as mean and ± s.d.

Type A spheroids, i.e., those formed during the 24 hours after the inoculation of shed cells into the agarose 600 μm pin molds, presented a spherical morphology with well-grouped cells forming a small and compacted spheroid (Fig. 1A).

Type B spheroids, i.e., those formed early in the 24 hours in the 600 μm agarose pin molds and then transferred to the periapical lesion molds, presented a compact and spindle-shaped morphology (Fig. 1B).

Type C spheroids from shed cells cultured in 2D monolayers and transferred to the periapical lesion mold showed a spherical morphology comparable to the type A spheroids but with a larger size. In this type of spheroid, we observed a hollow nucleus without cells (Fig. 1C).

Three spheroid growth curves were constructed according to the initial inoculum of cells and the type of agarose mold (600 μm pin mold or periapical lesion mold). In all three types of curves, we observed the growth phase, the stationary phase, and the decline phase, with this last phase having proportionally more dead than living cells.

For type A spheroids, i.e., when the shed cells from 2D monolayer cultures were seeded in 600 μm molds, the growth of the spheroids occurred between 24 and 48 hours (Fig. 1D). For type B spheroids, i.e., when preformed spheroids (early 24 hour spheroids formed in 600 μm molds) were transferred to the periapical lesion molds, their growth occurred immediately from the inoculation time to 24 hours after (Fig. 1E), and for type C spheroids, i.e., when the shed cells were seeded in periapical lesion molds, the growth of the spheroids occurred between 8 and 16 hours (Fig. 1F).

### Viability of the shed cells in monolayers and spheroids

The MTT test was performed to verify cell viability in cultures of shed cells with osteoblastic induction medium for 3, 7 and 10 days to check whether cell death of the cultures had occurred during this process. The results show that over the differentiation periods, the cultures of shed cells with DMEM/F12 medium supplemented with inducing agents showed an increase in viability (Fig. 2A).

**Figure 2.**
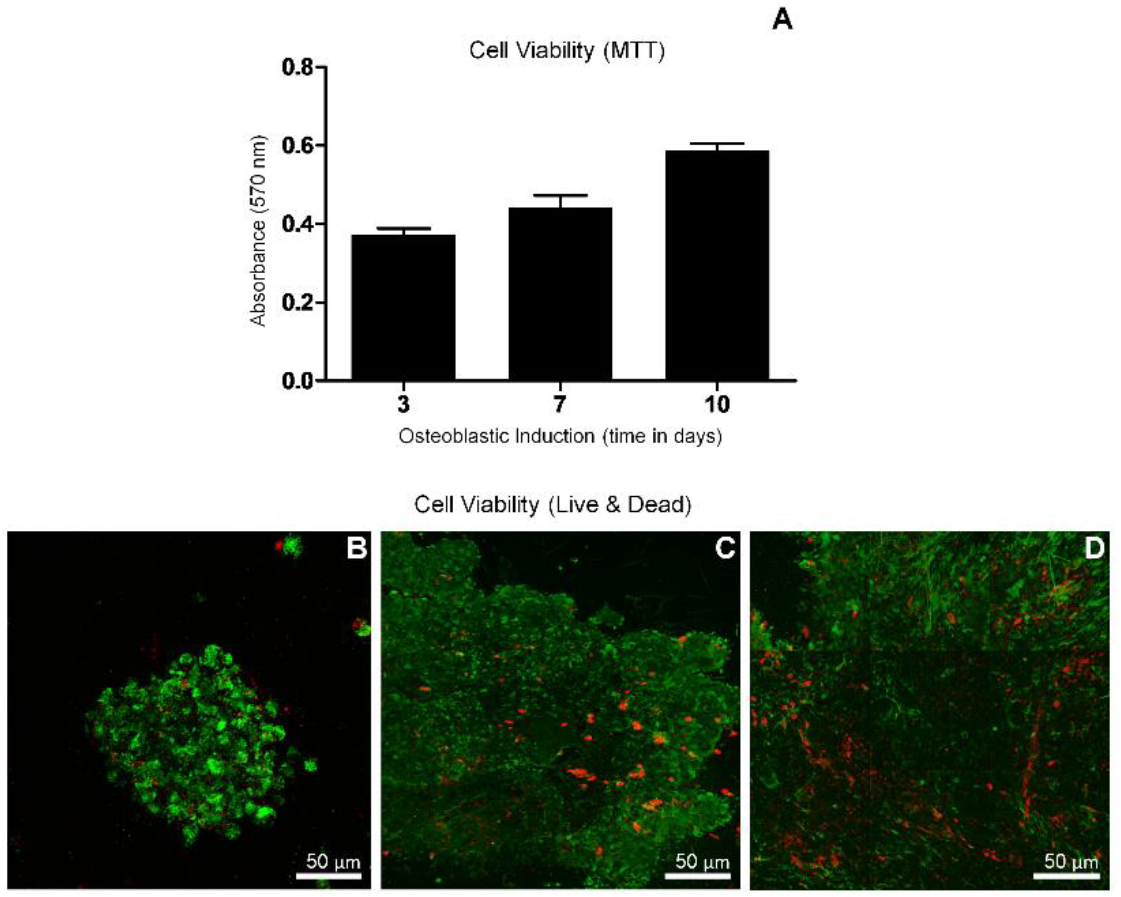
Viability of shed cells grown in monolayer and of spheroids. Shed cells were cultured in a 2D monolayer in the presence of an osteoblastic induction medium and observed after 3, 7, and 10 days. The MTT test showed that the viability of these cells was not negatively affected at the indicated times (Fig. 2A). For spheroids, viability was assessed using fluorescence microscopy (Live / Dead kit). Type “A” spheroids, grown for 24 hours (600 μm mold), showed a high rate of live cells (stained green) with some dead cells (stained red) (Fig. 2B). Type “B” spheroids, grown for 24 hours in 600 μm mold and then transferred to periapical lesion mold, are spindle-shaped and have many alive and some dead cells (Fig. 2C). Type “C” spheroids, shed cells grown in monolayer and then transferred to periapical lesion mold, also showed most living cells, but with the central region composed of dead cells (Fig.2D).

The viability of type A spheroids collected after 24 hours of development was shown to be high, as detected by the live/dead test by fluorescence microscopy. We observed that these spheroids showed spherical morphology with a large number of live cells stained green throughout their length but with the presence of a few apoptotic cells stained red (Fig. 2B).

Type B spheroids were predominantly fusiform with mostly viable cells, with only a few apoptotic cells dispersed in the periphery (Fig. 2C).

Type C spheroids showed spheroidal morphology with most living cells on the periphery and a cluster of apoptotic cells in the central region, typical of multicellular spheroids (Fig. 2D).

### Osteoblastic differentiation of the shed cells in monolayers and spheroids

Due to the importance of alkaline phosphatase (ALP) in mineralization and bone formation processes, ALP protein expression measurements were performed using the Fast Red technique.

The measurements were taken at 7, 10, 14, and 21 days of differentiation (n = 5) in the shed cells in the 2D monolayer culture with medium plus osteoblastic differentiation-inducing agents.

We observed a significant increase in ALP activity on days 10 and 14 and a decline on day 21 during *in vitro* osteoblastic differentiation of shed cells (Fig. 3A-B).

**Figure 3.**
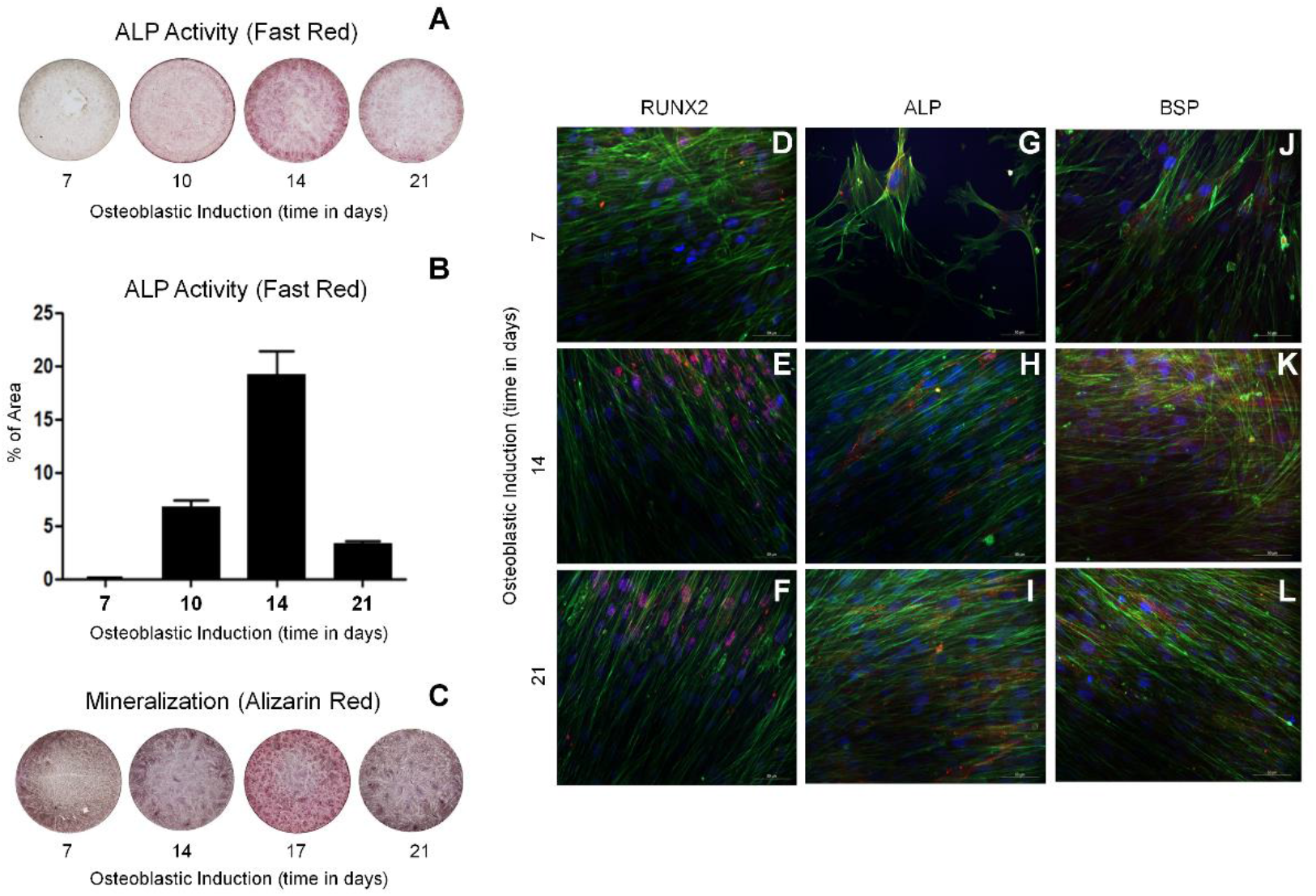
Osteoblastic differentiation of shed cells in monolayer. Shed cells were cultured in the presence of osteoblastic differentiation medium. The fast-red technique quantified the activity of alkaline phosphatase (ALP) in 7 to 21 days. The ALP activity peak occurred at 14 days, with the most significant decline at 21 days in culture (Figs 3A and 3B). Mineralization was also evaluated during differentiation by staining with alizarin red. The highest concentration of mineralization nodules occurred between 17 and 21 days in culture (Fig. 3C). Immunolocalization of differentiation markers RUNX2, ALP, and BSP was also used. Shed cells were cultured in the presence of an induction medium and observed after 7, 14, and 21 days. It was possible to follow from the 14 days in culture that the shed cells exhibited the three marker proteins (Figs. 3D-L).

The mineralization process was evaluated by the staining procedure with alizarin red, which allows qualitative data on the calcium deposits in the mineralization areas.

The stained shed cell cultures (n = 5) at 7, 14, 17, and 21 days during osteoblastic differentiation were imaged. The imaging analysis showed the presence of darkened areas as an indicator of mineralization nodules between 17 and 21 days (Fig. 3C).

The immunolocalization of three osteoblastic differentiation markers, the transcription factor RUNX2, the ALP enzyme, and bone sialoprotein (BSP), was performed during 7, 14, and 21 days of differentiation of shed cells in the 2D monolayer cultures. RUNX2 and ALP were expressed during the three analyzed periods, with RUNX2 located in the nucleus (Fig. 3D-F), ALP on the cell surface (Fig.3G-I), and BSP in the osteoblast cytoplasm (Fig.3J-L). All markers showed the same profile with increased expression at 21 days of differentiation.

To evaluate the relative transcriptional expression levels of osteoblastic differentiation markers, such as Runx2, Alp, Bglap, Sp7 and Bsp, in shed cells cultured in 2D monolayers or in 3D molds for spheroid formation during osteoblastic differentiation from 7 to 14 days, we performed RT-qPCR assays. Runx2 mRNA was significantly differentially expressed in both types of samples (shed cells grown in 2D monolayers or in 3D spheroids formed in periapical lesion molds) (Fig. 4A). Alp mRNA was differentially expressed when 3D spheroids were formed either in the 600 μm or in periapical lesion molds (Fig. 4B). Bglap mRNA was differentially expressed in 3D spheroids formed in the 600 μm molds (Fig. 4C). SP7 was differentially expressed as much in the shed cells cultured in 2D as in the 3D spheroids formed in molds of the periapical lesion (Fig. 4D), and although Bsp mRNA was expressed, its profile was not different between the tested samples (Fig. 4E).

**Figure 4.**
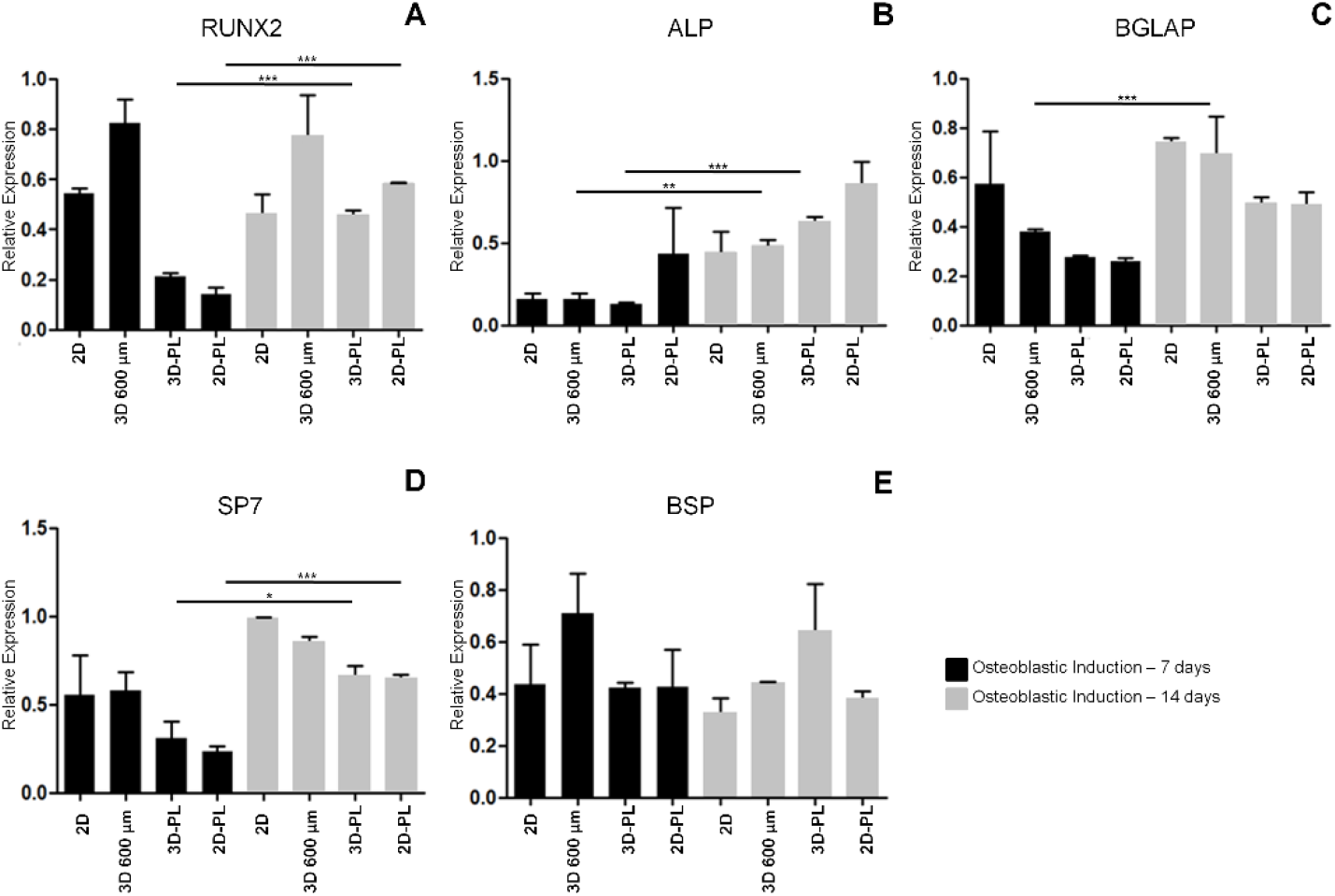
Relative expression of the transcripts that encode osteoblastic differentiation markers (RUNX2, ALP, BGLAP, SP7 and BSP) in the samples: shed cells grown in 2D monolayer culture (2D), spheroids grown in 600 μm molds (3D 600 μm), spheroids grown in 600 μm and transferred to periapical lesion molds (3D-PL) and shed cells grown in 2D monolayer culture and transferred to periapical lesion molds (2D-PL) for 7 and 14 days. GAPDH was used as housekeeping expression. The difference between the groups was assessed using the *t* statistical test. *** p <0.001; ** p <0.01; * p < 0.05. The appropriate independent triplicates were performed.

### Transcriptional expression of mesenchymal stemness markers

Mesenchymal stemness markers, such as Cd13, Cd29, Cd44, Cd73, and Cd90, mRNAs were also tested. We observed that the transcriptional expression of these markers, as a whole in the different samples (shed cells cultured in 2D monolayers and spheroids formed either in 600 μm or in periapical lesion molds), decreased significantly during osteoblastic differentiation (Supp. Figs 1A-E).

## Discussion

The persistence, recurrence, and even refractoriness of patients to synthetic materials used in the treatment of periapical lesions represent important challenges for regenerative endodontics. Hence, we asked whether spheroids originating from shed cells could be used in the biological restoration of such lesions, as previously considered (34).

The causes of these complications may be due to the root canal system’s complexity, leakage of the filling materials, extra-root infections, and actual cystic lesions (35,36). The host’s intrinsic factors also have to be taken into account, such as the body’s immune response against repairing the periapical lesion (6,37).

Genetic polymorphisms have been associated with adverse reactions against endodontic treatment, such as the imbalance in the expression of genes involved in the healing process, with consequences for the persistence or recurrence of the lesion (6,38).

Our objective was to establish an *in vitro* model system to test the biological recomposition of a periapical lesion cavity using shed cells. We compared the behavior of these cells grown in monolayers (2D) and spheroids (3D) in an artificial model assembled by 3D printing, which mimics a periapical lesion of an upper lateral incisor tooth.

Shed cells are promising for use in bone regeneration due to their low immunogenicity, high proliferative rate, and responsiveness to differentiation inducers (8,17,39).

We started from a micro tomographic image of a periapical lesion of an upper lateral incisor tooth. This image was used to make a negative impression of the lesion in acrylic using a 3D printer. This mold was used to prepare a positive mold of the lesion in agarose filled with shed cells from monolayer cultures or with spheroids.

During the formation of tissues *in vivo*, the cells interact with each other through extracellular matrix components and assume a 3D organization (40,41). We used spheroids to fill the periapical lesion mold and followed its differentiation into osteoblasts.

The spheroids of shed cells presented a spherical morphology with high viability. The growth curves demonstrated the exponential, stationary, and decline phases, as expected.

After being inoculated in the periapical lesion mold, we observed that in less than 24 hours, the spheroids started to aggregate in a spheroid-spheroid format, finally generating larger spheroids with different shapes and with high viability. This property might be important when using spheroids in the endodontic treatment of periapical lesions.

The culture of shed cells in a 2D monolayer inoculated directly into the periapical lesion mold generated single large spheroids formed in less than eight hours. In these spheroids, viable cells were mainly located on the surface, and apoptotic cells were located in the central zone.

This is a characteristic seen in large spheroids, which can form a hollow vascular lumen-like center through the apoptosis of polarized central cells, forming a layer of dead cells, resulting from hypoxia due to the difficulty of oxygen delivery. The nutrition of these cells located in the central spheroid area is also hampered (41,42), which may have caused a drop in cell viability compared with the viability of those preformed smaller spheroids, and both types were inoculated into the periapical lesion mold.

Then, we tested the osteoblastic differentiation of shed cells and spheroids. When subjected to differentiation, the shed cells from monolayer cultures proliferated and showed an increase in viability.

The activity of alkaline phosphatase (ALP) protein increased in the initial periods, decreasing in the later stages of differentiation. This was consistent since ALP represents a marker of the early stages of osteoblastic differentiation (43,44). The mineralization process and the immunolocalization of the RUNX2, ALP, and BSP protein markers proved to be prominent in the late phase of differentiation. This showed that the shed cell *in vitro* differentiation model system was adequate.

Analysis of the transcriptional expression of the respective mRNAs encoding mesenchymal stem cell markers (stemness markers) Cd13, Cd12, Cd44, Cd73 and Cd90 showed that these genes temporally decreased their expression as osteoblastic differentiation progressed. Concomitantly, the expression of the differentiation markers Runx2, Alp, Bglap, and Sp7 mRNAs showed that they increased at the beginning of osteoblastic differentiation (7 to 14 days) in both the shed cells of the monolayer and spheroid cultures, except for Bsp mRNA, whose expression remained uniform during that period.

The biological restoration of periapical lesions represents an attractive candidate for endodontics. It might be possible to establish a clinical protocol that begins with spheroid implantation from shed cells at the beginning of osteoblastic differentiation. Over time and with the occurrence of mineralization, the lesion and the cells would be gradually replaced by bone tissue formed *in situ*, ruling out any adverse reactions from the host.

## Conclusion

The results of this study show that we were able to develop an *in vitro* model system that mimics a periapical lesion with the possibility of repopulating the lesion cavity with shed cell spheroids. Additionally, the model made it possible to differentiate spheroid cells into osteoblasts that mineralize over time. This opens up a new perspective for endodontics for restoring areas with periapical lesions through replacement with *in situ* bone tissue formation.

## Acknowledgments

This work was partially funded by the National Council for Scientific and Technological Development (CNPq, Brasília, Brazil, through the grant No. 305787/2017-9) and CAPES (Brasília, Brazil, through the financial code 001). Our laboratories are funded by São Paulo Research Foundation (FAPESP, São Paulo, Brazil). We thank Dr. Márcio M. Beloti, Dr. Paulo Tambasco de Oliveira, Dr. Selma Siéssere, Dr. Adalberto L. Rosa, and Dr. Ricardo Gariba da Silva from the School of Dentistry of Ribeirão Preto, USP, Ribeirão Preto, Brazil for their help and discussions.

---

**Supplemental Figure 1.**
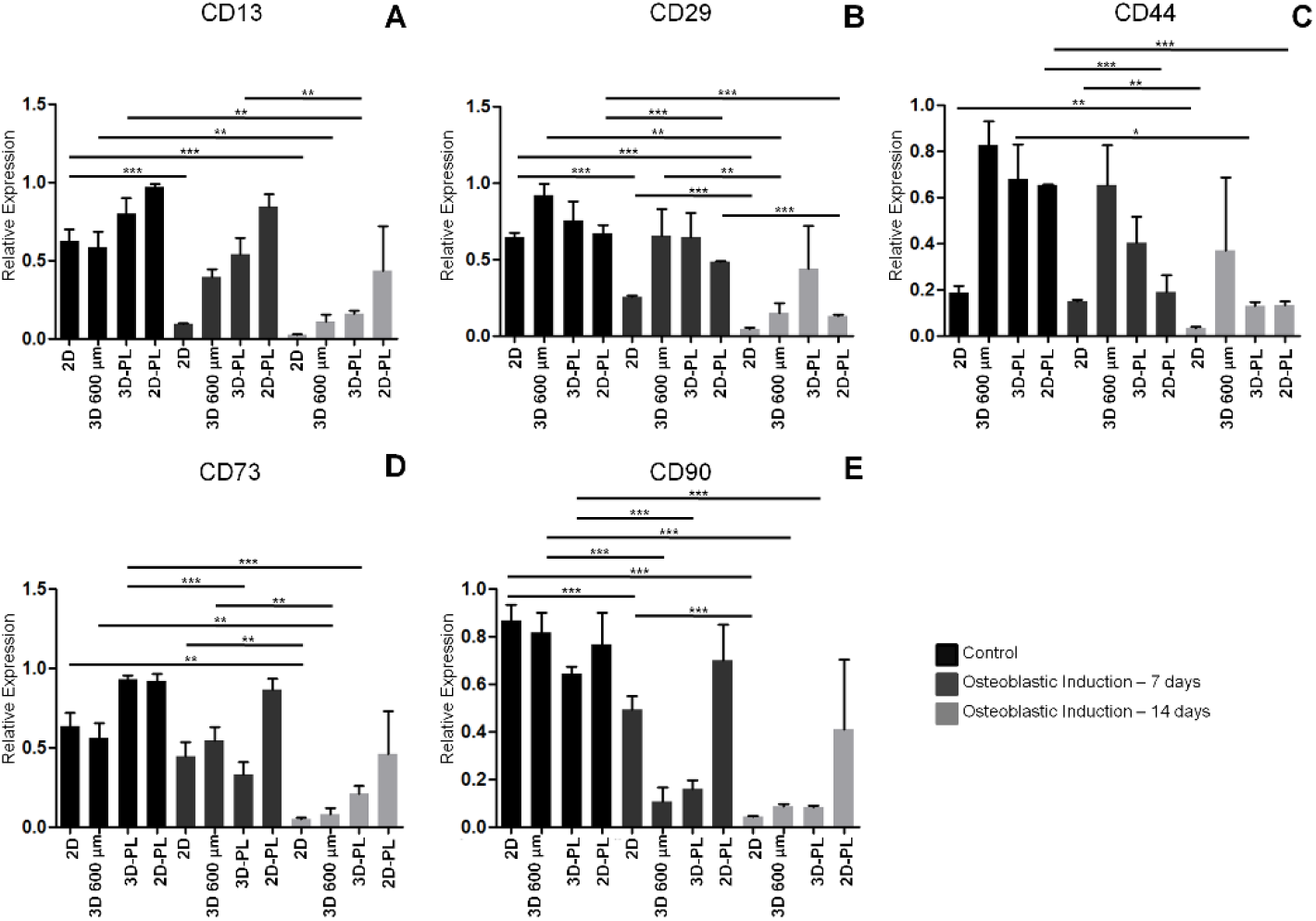
Relative expression of the transcripts that encode stemness markers (Cd13, Cd29, Cd44, Cd73 and Cd90) in the samples: shed cells grown in 2D monolayer culture (2D), spheroids grown in 600 μm molds (3D 600 μm), spheroids grown in 600 μm and transferred to periapical lesion molds (3D-PL) and shed cells grown in 2D monolayer culture and transferred to periapical lesion molds (2D-PL) comparing undifferentiated (control) with osteoblastic differentiated cells for 7 and 14 days. GAPDH was used as housekeeping expression. The difference between the groups was assessed using the ANOVA One-Way test. *** p <0.001; ** p <0.01; * p <0.05. The appropriate independent triplicates were performed.

